# Anxiety modulates perception of facial fear in a pathway-specific, lateralized, manner

**DOI:** 10.1101/141838

**Authors:** Hee Yeon Im, Reginald B. Adams, Jasmine Boshyan, Noreen Ward, Cody A. Cushing, Kestutis Kveraga

## Abstract

Facial expression and eye gaze provide a shared signal about threats. While averted-gaze fear clearly points to the source of threat, direct-gaze fear renders the source of threat ambiguous. Dual processing routes have been proposed to mediate these processes: reflexive processing via magnocellular (M-) pathway and reflective processing via parvocellular (P-) pathway. We investigated how observers’ trait anxiety modulates Mand P-pathway processing of clear and ambiguous threat cues. We performed fMRI on a large cohort (N=108) widely ranging in trait anxiety while they viewed fearful or neutral faces with averted or directed gaze. We adjusted luminance and color of the stimuli to selectively engage M- or P-pathway processing. We found that higher anxiety *facilitated* processing of averted-gaze fear projected to M-pathway, but *impaired* perception of direct-gaze fear projected to P-pathway. Increased right amygdala reactivity was associated with higher anxiety, only for averted-gaze fear presented to M-pathway. Conversely, increased left amygdala reactivity was associated with higher anxiety for P-biased, direct-gaze fear. This lateralization was more pronounced with higher anxiety. Our findings suggest that trait anxiety has differential effects on perception of clear and ambiguous facial threat cues via selective engagement of M and P pathways and lateralization of amygdala reactivity.

## Introduction

Facial expression and direction of eye gaze are two important sources of social information. Reading emotional expression can be informative in understanding and forecasting an expresser’s behavioral intentions [1–5], and understanding eye gaze allows an observer to orient spatial attention in the direction signaled by the gaze [6–11]. Individuals with many cognitive disorders show distorted ability in perceiving social cues from faces (e.g., generalized anxiety disorder [12,13], social anxiety disorder [14,15], and depression [16] or impaired (e.g., prosopagnosia [17,18] and autism [19,20]).

The signals that facial expression and eye gaze convey interact in the perceiver’s mind. A fearful facial expression with an averted gaze is typically recognized as indicating a threat located where the face is “pointing with the eyes” [21]. Both fear and averted gaze are avoidance-oriented signals and together represent congruent cues of threat, where the eye gaze direction indicates the potential location of the threat causing the fear in the expresser [22–26]. Conversely, direct gaze is an approach-oriented cue directed at the observer, while fear is an avoidance cue. Thus, unless the observer is the source of threat, fear with direct gaze is a more ambiguous combination of threat cues because it is unclear whether the expresser is signaling danger or attempting to evoke empathy [21]. Consistent with these interpretations, observers tend to perceive averted-gaze fear as more intense and recognize it more quickly and accurately, compared to direct-gaze fear [1,24–28]. Eye gaze also influences the perception of emotion in neutral faces: Approach-oriented emotions (anger and happy) are attributed to neutral faces posed with direct gaze whereas avoidant-oriented emotions (fear and sadness) are attributed to neutral faces posed with averted gaze [25,29].

Neuroimaging studies have highlighted the role of the amygdala in such integration of emotional expression with eye gaze [21,23,25,26,30–36]. The amygdala reactivity to the interaction of facial expression with eye gaze has been shown to be modulated by presentation speed [23,30,33] and this modulation differs by hemisphere [23,30,33,37]. These findings suggest that the bilateral amygdalae are differentially involved in processing of clear, congruent threat cues (averted-gaze fear) and ambiguous threat cues (direct-gaze fear). Specifically, left and right amygdalae showed heightened activation to longer exposures of direct-gaze fearful faces (ambiguous threat), and to shorter exposures of averted-gaze fearful faces (clear threat), respectively [23,30,33]. This suggests that the right amygdala may be more involved in early detection of clear threat cues and the left amygdala may engage more in slow reassessment and evaluation of ambiguous threat cues.

According to dual process models [38–41], “reflexive” and “reflective” processes operate as a fast, automatic response, and a relatively effortful, top-down controlled process for finer-tuned information processing, respectively. The dual process model also has a direct parallel in the threat perception literature, which has proposed the existence of the so-called “low road” vs. “high road” routes [42–44]. The “low-road” is considered to be an evolutionarily older pathway (presumably via the coarser, achromatic magnocellular (M) pathway projections), for processing of “gist” and rapid defensive responses to threat without conscious thought [42,45–47]. The “high-road”, on the other hand, is considered as the route for a slower, conscious processing of detailed information, allowing for modulation of initial low-road processing [42,43,45] and might be subserved predominantly by the parvocellular (P) pathway projection through the ventral temporal lobe.

Current models of face perception also support similar, parallel pathways in the human visual system (M and P) that are potentially tuned to different processing demands. For example, Vuilleumier et al. [48] suggested that low spatial frequency (LSF) and high spatial frequency (HSF) information of fearful face stimuli are carried in parallel via M- and P-pathways, respectively. By exploiting face stimuli designed to selectively bias processing toward M- vs. P-pathway, Adams et al. [31] recently found brain activations for averted-gaze fear faces (clear threat) in M-pathway regions and for direct-gaze fear faces (ambiguous threat) in P-pathway regions. Such dissociation in the responsivity of the Mand P-pathways to clear and to ambiguous threat signals, respectively, led the authors to suggest that fearful face by eye gaze interaction may engage a similar, generalizable dual process of threat perception that engages reflexive (via M-pathway) and reflective (via P-pathway) responses [31].

Although substantial individual differences in behavioral responses and amygdala reactivity to emotional faces have been closely associated with observers’ anxiety [13,49–54], only a few studies have systematically examined the role of anxiety in such integrative processing of facial fear and eye gaze [11,29,55–57]. They found that high trait-anxiety individuals show more integrative processing of facial expressions and eye gaze for visual attention [11], stronger cueing effect by eye gaze in fearful expressions [56,57], and increased amygdala reactivity to compound threat-gaze cues [29], suggesting that high anxiety level is associated with higher sensitivity and stronger reactivity to compound threat cues in general. To date, none of the studies has examined how anxiety differentially modulates amygdala reactivity to clear and ambiguous threat cues that are preferentially conveyed via different (M and P) visual pathways. Thus, better understanding of how perceivers’ anxiety modulates the processing of facial threat cues would contribute to the refinement of behavioral and neuroanatomic models for anxiety-related disorders.

The primary aim of the current study was to examine how perceiver’s anxiety modulates behavioral and neural responses to averted-gaze fear (clear threat) vs. direct-gaze fear (ambiguous threat) in a larger (N of 108) and more representative cohort, compared to previous studies that exploited relatively modest sample sizes (e.g., N=27 [49], N=31 [29], and N=32 [54]). Based on the recent findings [31] that compound threat cues differentially favor visual input via M- and P-pathways depending on the clarity or ambiguity of threat, we hypothesized that the association between high anxiety and increased amygdala reactivity would be observed specifically for averted-gaze fear faces projected to M-pathway, and for direct-gaze fear projected to P-pathway.

Furthermore, we also wanted to examine how anxiety-related modulation of neural activity varies between the hemispheres, given the previous evidence for hemispheric difference in the amygdala activity in perception of facial fear [23,31,33,58,59]. More right amygdala involvement was observed in responding to subliminal threat stimuli [59] and briefly-presented, averted-gaze fear faces (clear threat cues: [23,33]) or averted-gaze eyes [58]; whereas more left amygdala involvement was observed in responding to supraliminal threat stimuli [59] and ambiguous threat cues conveyed by direct-gaze fear faces [23,31,58]. Therefore, another aim of our study was to directly test laterality effects in the activation pattern of the left and right amygdala in response to facial fear and their modulation by perceiver anxiety.

## Results

Participants (N=108) viewed images of fearful or neutral faces with direct and averted gaze with one-second presentations while undergoing fMRI. The stimuli were two-tone images of faces presented as high-luminance contrast (Unbiased), low-luminance contrast (M-biased), or isoluminant red/green, chromatically defined (P-biased) images. Participants were asked to report whether the face presented in the stimulus looked fearful or neutral. In order to investigate the effects of anxiety on the processing of compound facial cues presented in the magnocellular (M) and parvocellular (P) biased stimuli, we examined the relationship between participants’ anxiety and behavioral measurements (accuracy and RT) and amygdala activation during perception of these stimuli. We used trait anxiety rather than state anxiety to investigate the more enduring effects of anxiety [11]. Participants’ state anxiety scores ranged from 20 to 64 (mean=32.6, SD=9.4), and trait anxiety scores from 20 to 66 (mean=33.9, SD=9.7). The participants’ trait anxiety highly correlated with their state anxiety scores (*r* = 0.71, *p* < 0.001).

The beta estimates extracted from the amygdala and the behavioral measurements were screened for outliers (3 SD above the group mean) within each condition. As a result, 1.62% and 1.16% of the data points on average were excluded from response time (RT) and from accuracy, and 0.69% and 0.93% of the data points on average were excluded from the left and right amygdala activation for the further analyses. We report here the partial correlation coefficient *r* after controlling for the effects of the participants’ age and sex.

### Behavioral results

Figures 2A-2D show the mean response times (RT) of the correct trials and mean accuracy as a function of trait anxiety scores, when participants viewed the M- and P-biased stimuli containing fearful faces with averted or direct eye gaze. The participants with high trait anxiety made faster responses to averted-gaze fear than those low in trait anxiety, for both the M-biased and P-biased stimuli (Figure 2A, thicker regression lines indicate statistically significant trends), although only the RTs in the P-biased stimuli reached statistical significance after regressing out the participants’ age and sex (M-biased: *r* = −0.175, *p* = 0.070; P-biased: *r* = −0.229, *p* < 0.02). The median RT showed similar patterns: M-biased: *r* = −0.187, *p* = 0.054; P-biased: *r* = −0.208, *p* < 0.02. Furthermore, participants with higher-trait anxiety were more accurate in recognizing averted-gaze fear in the M-biased stimuli (Figure 2B: *r* = 0.218, *p* < 0.03), but not in the P-biased stimuli (*r* = 0.071, *p* = 0.470).

**Figure 1:**
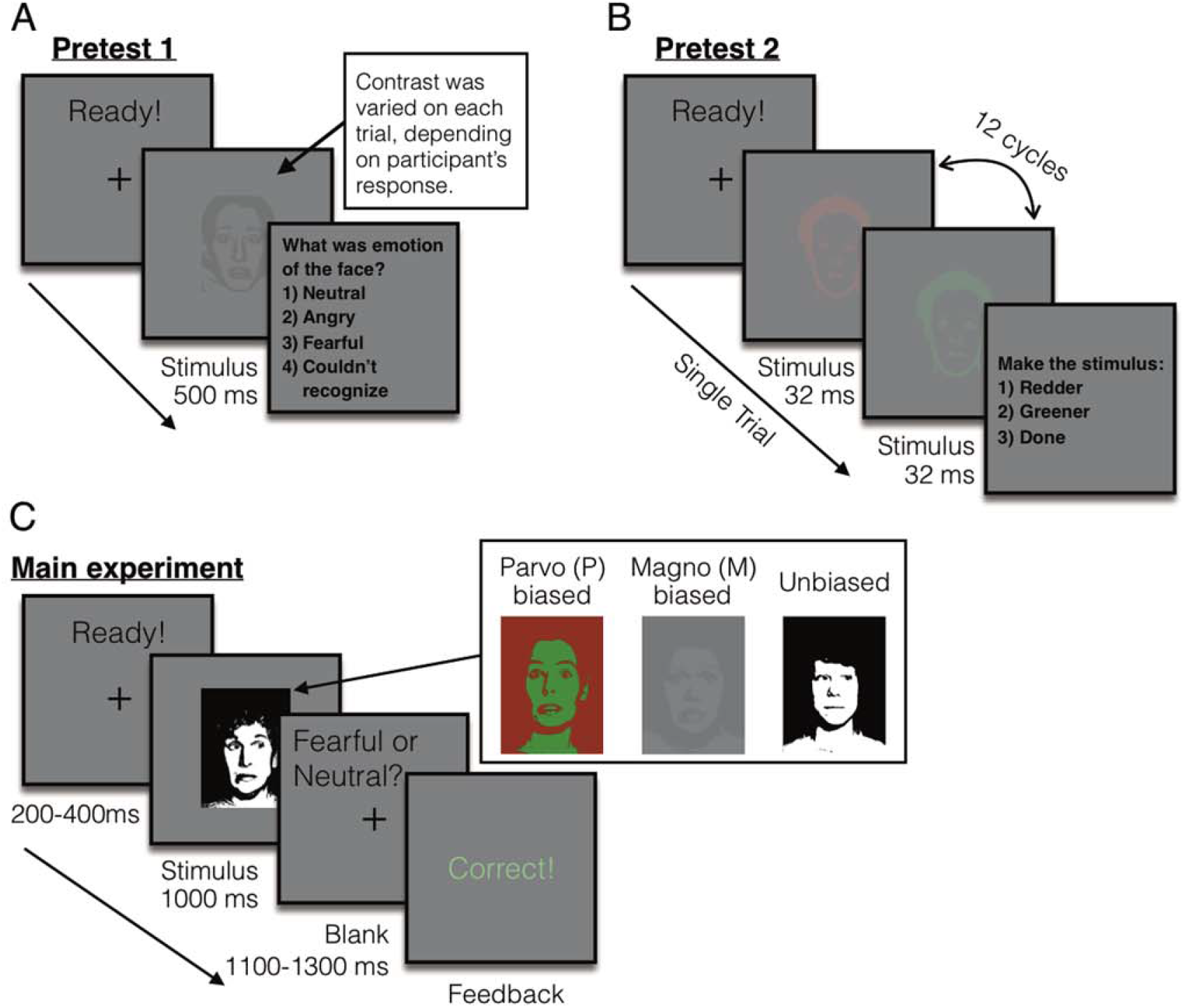
Sample trials of the pretests and the main experiment. (A) A sample trial of pretest 1 to measure the participants’ threshold for the foreground-background luminance contrast for achromatic M-biased stimuli. (B) A sample trial of pretest 2 to measure the participants’ threshold for the isoluminance values for chromatic P-biased stimuli. (C) A sample trial of the main experiment.

**Figure 2:**
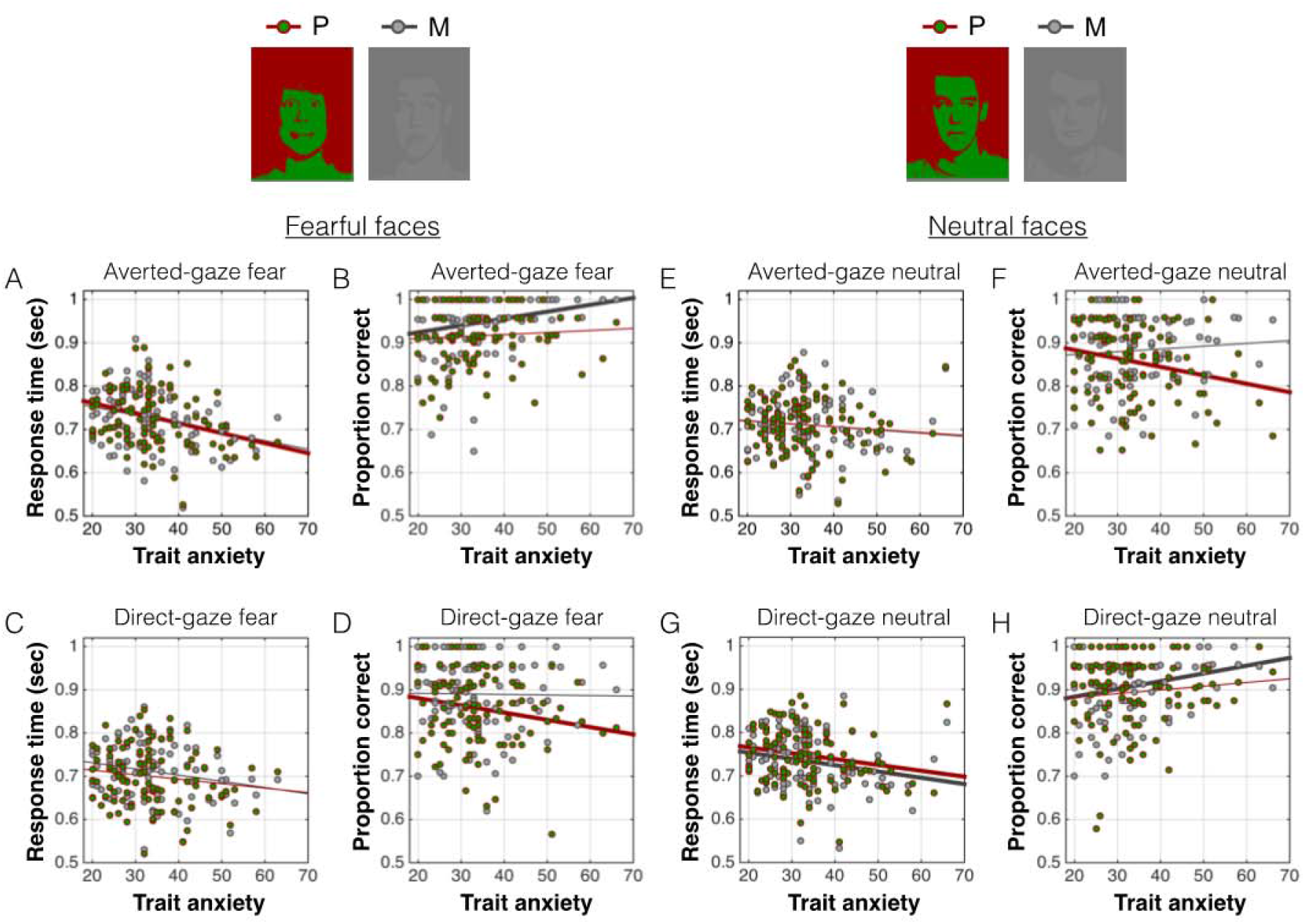
The behavioral results from the main experiment. (A) The response time (RT) for the fearful faces with averted eye gaze, presented in the M-biased (dots in gray) and in the P-biased (dots in red-green) stimuli. The gray and red lines indicate the linear relationship between trait anxiety and the RT for the M-biased stimuli and the P-biased stimuli, respectively. The thicker lines indicate statistically significant correlations. (B) The accuracy for the fearful faces with averted eye gaze, presented in the M-biased (dots in gray) and in the P-biased (dots in red-green) stimuli. The thicker lines indicate statistically significant correlations. (C) The RT for the fearful faces with direct eye gaze, presented in the M-biased (dots in gray) and in the P-biased (dots in red-green) stimuli. (D) The accuracy for the fearful faces with direct eye gaze, presented in the M-biased (dots in gray) and in the P-biased (dots in red-green) stimuli. The thicker lines indicate statistically significant correlations. (E-F) The RT and the accuracy for the neutral faces with averted eye gaze. (G-H) The RT and the accuracy for the neutral faces with direct eye gaze.

For the processing of direct-gaze fear, we did not observe significant trends between the trait anxiety and the mean RTs for the M-biased (Figure 2C: *r* = −0.106, *p* = 0.281) nor P-biased stimuli (*r* = −0.026, *p* = 0.790). The median RT showed similar patterns: M-biased: *r* = −0.147, *p* = 0.132; P-biased: *r* = −0.106, *p* < 0.278. As in Figure 2D, participants with high trait anxiety showed impaired accuracy for recognizing direct-gaze fear only when it was presented in the P-biased stimuli (*r* = −0.220, *p* < 0.03), but not in the M-biased stimuli (*r* = −0.010, *p* = 0.922).

We next examined the effects of the trait anxiety on the perception of neutral faces. Unlike the fearful faces, the participants’ mean RTs to neutral faces with averted eye gaze did not show significant trend in the M-biased (Figure 2E, *p* = 0.890) nor P-biased (Figure 2E, *p* = 0.993), and the same was true for the median RTs (*p’s* > 0.679). Participants with higher-trait anxiety made more error responses for neutral faces with averted eye gaze in the P-biased condition (Figure 2F, *r* = −0.233, *p* < 0.02), but not in the M-biased condition (*p* = 0.510). For direct-gaze neutral faces, the participants with higher trait anxiety showed faster mean RTs for both M- and P-biased stimuli (Figure 2G, M-biased: *r* = −0.229, *p* < 0.02; P-biased: *r* = −0.211, *p* < 0.03) and faster median RTs, as well (M-biased: *r* = −0.168, *p* = 0.084; P-biased: *r* = −0.164, *p* = 0.092). For the accuracy, we found the selective facilitation of the processing of M-biased neutral faces with direct eye gaze in the high trait anxiety participants (Figure 2H: *r* = 0.196, *p* < 0.05), but not the processing of P-biased stimuli (*p* = 0.438).

Therefore, the behavioral results show that when *clear* cue combinations are presented (averted-gaze fear or direct-gaze neutral), increased trait anxiety had the effect of reducing RTs for faces presented in both M- and P-biased form, but recognition accuracy became significantly higher only for the *M-biased* stimuli. Conversely, when *ambiguous* cue combinations (direct-gaze fear and averted-gaze neutral are presented, increased trait anxiety did not reduce the RTs, and recognition accuracy decreased, but only for the *P-biased* stimuli. Taken together, these results suggest that trait anxiety interacts with visual pathway processing biases, such that clear facial cue recognition is improved in the M-pathway, and ambiguous facial cue recognition is *impaired* in the P-pathway, with increased trait anxiety.

### fMRI results

Table 1 presents the full list of activations (threshold: *p* < 0.001, extent: k = 4) for each of the contrasts of our interest: M averted fear – M direct fear, M direct fear – M averted fear, P direct fear – P averted fear, and P averted fear – P direct fear. Previous studies have reported that the left and right amygdala showed differential preferences for clear threat cue (e.g., averted fear) vs. ambiguous threat cue (e.g., direct fear) and for M-biased vs. P-biased stimuli [23,31]. We replicated these findings by showing that the right amygdala was preferentially activated by M-biased averted fear whereas the left amygdala was preferentially activated by P-biased direct fear (Figure 3).

**Figure 3:**
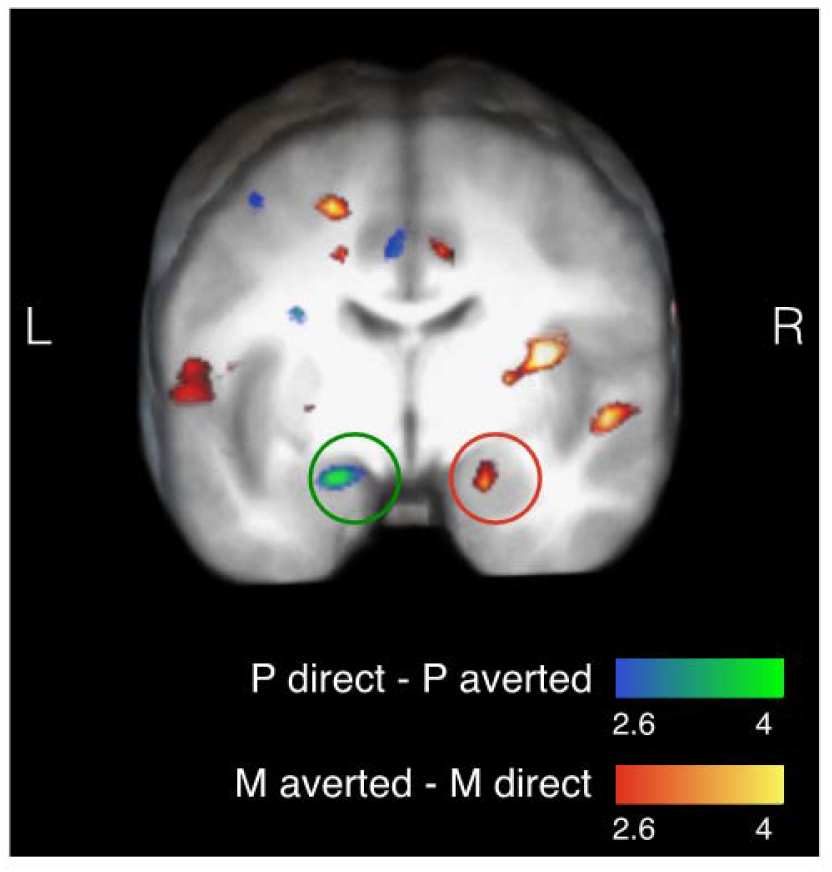
Activation map corresponds to whole-brain analyses showing the left and right amygdalae activations for P-biased direct fear minus P-biased averted fear and for M-biased averted fear minus M-biased direct fear, respectively.

**Table 1:**
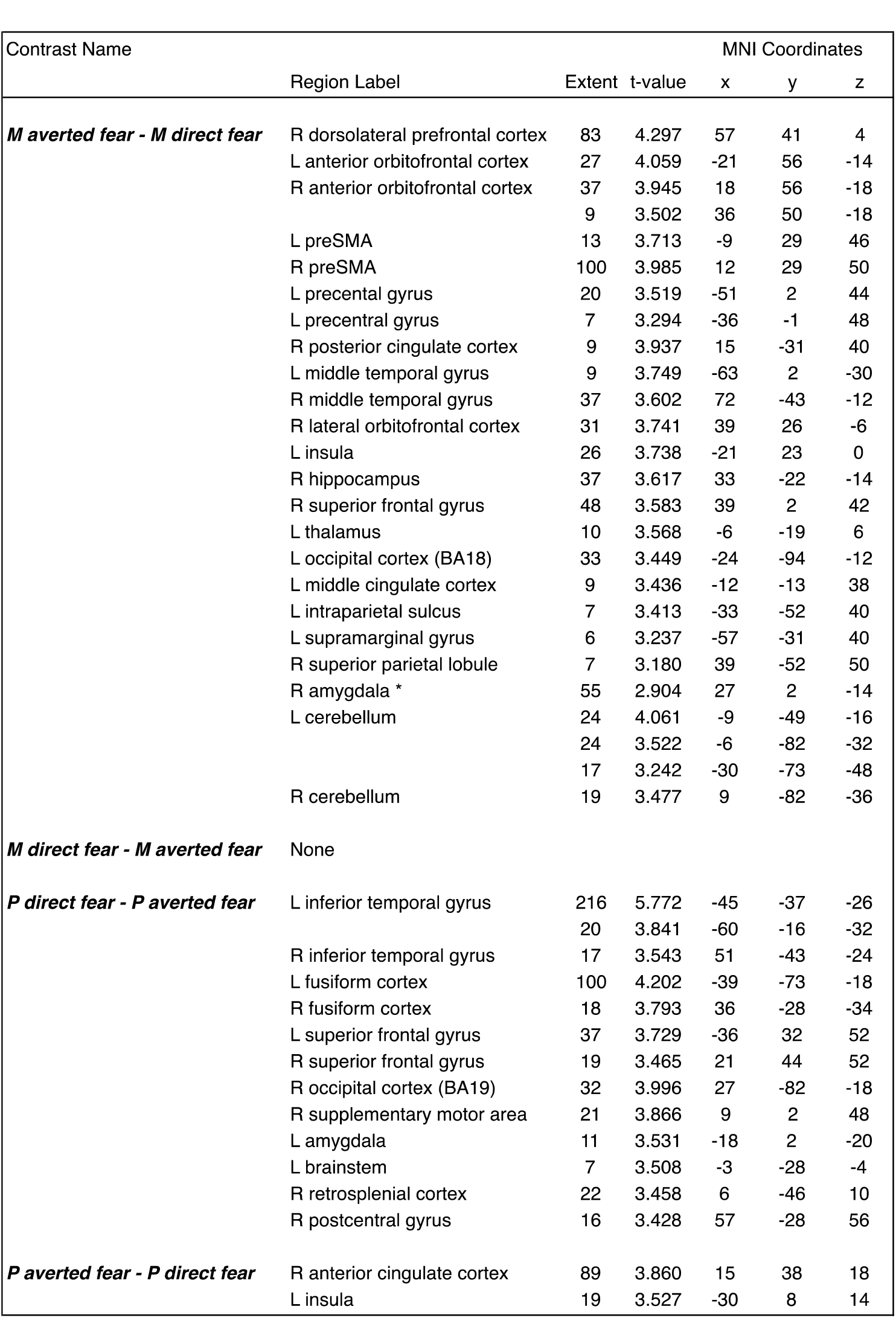
BOLD activations from group analysis, thresholded at *p* < 0.001 and *k* > 5, uncorrected. \**p* = 0.002.

We next examined the partial correlation coefficients (*r*) between the participants’ trait anxiety and the left and right amygdala activations when the participants viewed the Mand P-biased stimuli containing fearful and neutral faces with direct or averted eye gaze. As shown in Figure 4, we found that the observers’ trait anxiety also differentially modulated the left and right amygdala responses to averted-gaze fearful faces, presented in M-biased vs. P-biased condition. Although there was no significant trend between trait anxiety and the left amygdala reactivity to P-biased averted-gaze fearful faces (*r* = 0.091, *p* = 0.355), we observed the marginally significant negative correlation between trait anxiety and the left amygdala reactivity to M-biased averted-gaze fearful faces (Figure 4A; *r* = −0.174, *p* = 0.075). Moreover, we found that the right amygdala responses to M-biased averted-gaze fearful faces positively correlated with the trait anxiety (Figure 4B: *r* = 0.234, *p* < 0.02), even though there was no correlation of right amygdala responses with P-biased averted-gaze fearful faces: *r* = −0.010, *p* = 0.324).

**Figure 4:**
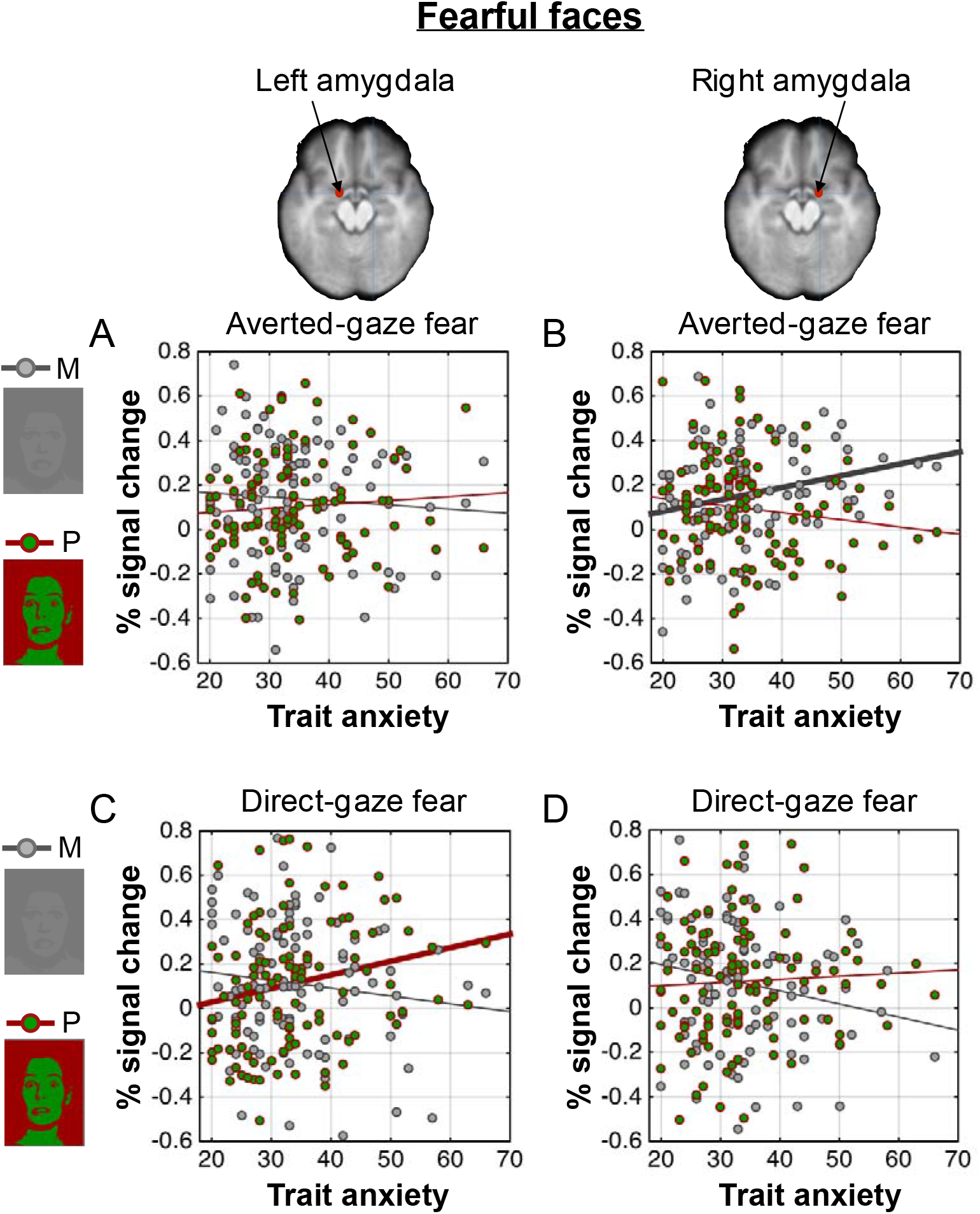
The left and right amygdala activation during perception of fearful faces. (A) The scatter plot of the trait anxiety and the % signal change in the left amygdala when participants viewed fearful faces with averted eye gaze presented in the M-biased (in gray dots) and in the P-biased (in red-green dots) stimuli. The gray and red lines indicate the linear relationship between trait anxiety and the left amygdala activation for the M-biased stimuli and the P-biased stimuli, respectively. The thicker lines indicate statistically significant correlations. (B) The scatter plot of the trait anxiety and the % signal change in the right amygdala when participants viewed fearful faces with averted eye gaze presented in the M-biased (in gray dots) and in the P-biased (in red-green dots) stimuli. The thicker lines indicate statistically significant correlations. (C) The scatter plot of the trait anxiety and the % signal change in the left amygdala when participants viewed fearful faces with direct eye gaze presented in the M-biased (in gray dots) and in the P-biased (in red-green dots) stimuli. (D) The scatter plot of the trait anxiety and the % signal change in the left amygdala when participants viewed fearful faces with direct eye gaze presented in the M-biased (in gray dots) and in the P-biased (in red-green dots) stimuli.

In response to direct-gaze fear faces, activity in the left amygdala showed significant, positive correlation with trait anxiety only when stimuli were presented in the P-biased form (Figure 4C; *r* = 0.196, *p* < 0.05), but there was no significant correlation for the M-biased stimuli (*r* = −0.125, *p* = 0.198). The right amygdala activation did not show significant trends for M-biased direct-gaze fear (Figure 4D: *r* = −0.119, *p* = 0.228) or for P-biased direct fear (*r* = −0.007, *p* = 0.947). These findings suggest that modulation by observers’ trait anxiety was selective depending on the emotional valence, eye gaze, and pathway biases, highly lateralized in amygdala activation: High trait anxiety was associated with the increased right amygdala activation for M-biased averted-gaze fearful faces (clear threat) and the increased left amygdala activation for P-biased direct-gaze fearful faces (ambiguous threat). Unlike for fearful faces, however, we did not observe any significant correlations between trait anxiety and the amygdala responses to neutral faces (Figure 5; all *p’s* > 0.370).

**Figure 5:**
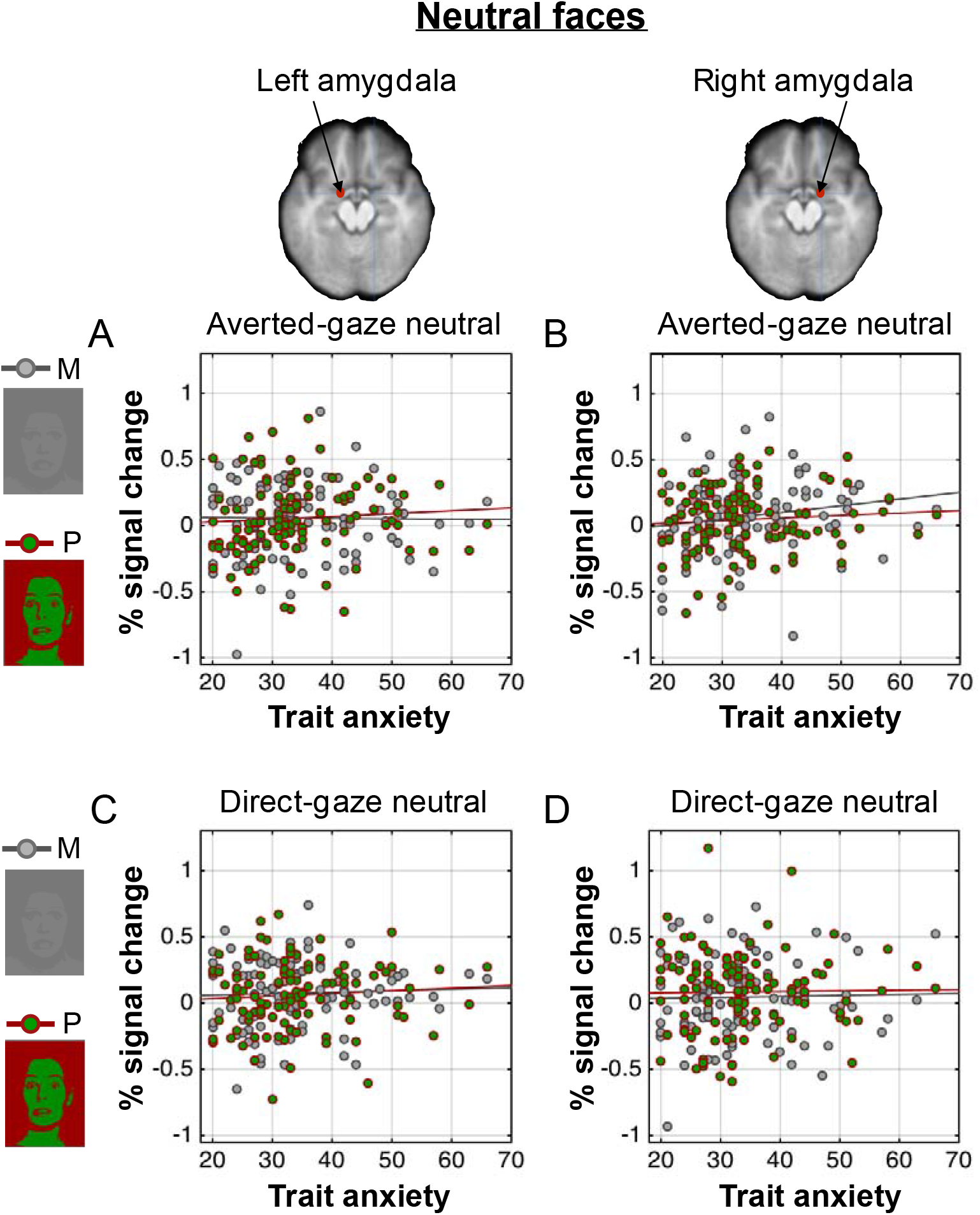
The left and right amygdala activation during perception of neutral faces. (A) The scatter plot of the trait anxiety and the % signal change in the left amygdala when participants viewed neutral faces with averted eye gaze presented in the M-biased (dots in gray) and in the P-biased (dots in red-green) stimuli. The gray and red lines indicate the linear relationship between trait anxiety and the left amygdala activation for the M-biased stimuli and the P-biased stimuli, respectively. (B) The scatter plot of the trait anxiety and the % signal change in the right amygdala when participants viewed neutral faces with averted eye gaze presented in the M-biased (dots in gray) and in the P-biased (dots in red-green) stimuli. (C) The scatter plot of the trait anxiety and the % signal change in the left amygdala when participants viewed neutral faces with direct eye gaze presented in the M-biased (dots in gray) and in the P-biased (dots in red-green) stimuli. (D) The scatter plot of the trait anxiety and the % signal change in the right amygdala when participants viewed neutral faces with direct eye gaze presented in the M-biased (dots in gray) and in the P-biased (dots in red-green) stimuli.

Because previous studies have suggested differential preference of the left and right amygdalae for P-biased direct fear and M-biased averted fear [23,31], we examined the magnitude of the hemispheric asymmetry by subtracting the left amygdala activation from the right amygdala activation for each condition. We plotted the difference between the activation levels in the left and the right amygdala as a function of the participants’ trait anxiety (Figure 6). Positive values indicate right amygdala activation greater than the left amygdala (e.g., right hemisphere (RH) dominant), whereas negative values indicate left amygdala activation greater than the right amygdala (e.g., left hemisphere (LH) dominant). While we did not observe significant lateralization for the neutral faces (all *p’s* > 0.198, Figures 6A and 6B), we found the significant lateralization for the fearful faces, systematically modulated by trait anxiety. The right amygdala became more dominant over the left amygdala with increasing trait anxiety, but only for the M-biased averted-gaze fear (Figure 6C; *r* = 0.289, *p* < 0.005), and not for P-biased averted-gaze fear (*r* = 0.095, *p* = 0.339). Conversely, the left amygdala became more dominant over the right amygdala with the increasing trait anxiety for the P-biased direct-gaze fear (Figure 6D; *r* = −0.236, *p* < 0.02), but not for the M-biased direct-gaze fear stimuli (*r* = −0.033, *p* = 0.740). These results suggest that observers with higher trait anxiety show more pronounced hemispheric lateralization between the left and right amygdala, with the right amygdala being more dominant for the processing of M-biased averted-gaze fear (clear threat) and the left amygdala being more dominant for the processing of P-biased direct-gaze fear (ambiguous threat).

**Figure 6:**
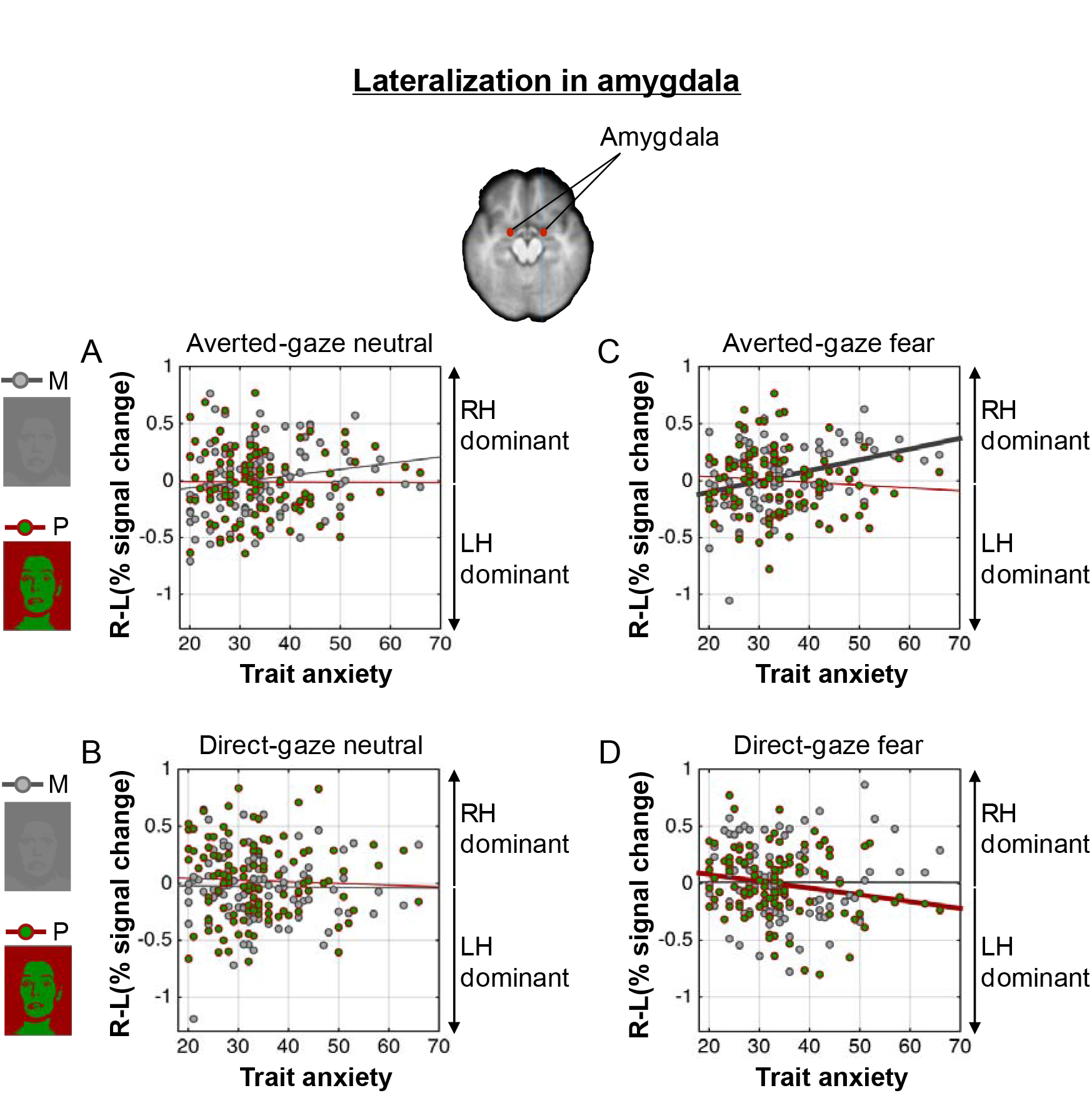
The magnitude of hemispheric lateralization in amygdala, obtained by subtracting the % signal change in the left amygdala from that in the right amygdala for each participant. The magnitude of lateralization is then plotted as a function of participants’ trait anxiety. The positive values indicate greater activation in the right amygdala (right hemisphere (RH) dominant) whereas the negative values indicate greater activation in the left amygdala (left hemisphere (LH) dominant). (A) The magnitude of hemispheric lateralization in amygdala when participants viewed neutral faces with averted eye gaze, presented in the M-biased (dots in gray) and in the P-biased (dots in red-green). The lines indicate the linear regressions with trait anxiety. (B) The magnitude of hemispheric lateralization in amygdala when participants viewed neutral faces with direct eye gaze, presented in the M-biased (dots in gray) and in the P-biased (dots in red-green). (C) The magnitude of hemispheric lateralization in amygdala when participants viewed fearful faces with averted eye gaze, presented in the M-biased (dots in gray) and in the P-biased (dots in red-green). The thicker regression lines indicate statistically significant correlations (D) The magnitude of hemispheric lateralization in amygdala when participants viewed fearful faces with direct eye gaze, presented in the M-biased (dots in gray) and in the P-biased (dots in red-green).

## Discussion

Anxiety can be hugely disruptive to everyday life, and over a quarter of the population suffers from an anxiety disorder during their lifetime [60]. The primary goal of this study was to examine how individuals’ trait anxiety modulates behavioral and amygdala responses in reading compound facial threat cues - one of the most common and important social stimuli-via two major visual streams, the magnocellular and parvocellular pathways. Here we employed a larger (N=108) and more representative (age range 20–70) community sample than in most of the previous studies examining the effects of anxiety on the affective facial processing [29,49,54]. We reported three main findings: 1) Compared to low trait-anxiety individuals, high trait-anxiety individuals were more accurate for M-biased averted-gaze fear (a clear threat cue combination), but less accurate for P-biased direct-gaze fear (ambiguous threat cues). The opposite was true for the neutral faces such that participants were better at correctly recognizing M-biased direct-gaze neutral faces, but less accurate for P-biased averted-gaze neutral faces; 2) increased right amygdala reactivity was associated with higher trait anxiety only for M-biased averted-gaze fear (clear threat), whereas increased left amygdala reactivity was associated with higher trait anxiety only for P-biased direct-gaze fear (ambiguous threat), and 3) the magnitude of this laterality effect for the fearful faces increased with higher trait anxiety. The current findings indicate differential effects of observers’ anxiety on threat perception as a result of the interplay of fearful face, eye gaze, and visual pathway processing biases.

Previous studies [12,61] and cognitive formulations of anxiety [62,63] have suggested that vulnerability to anxiety is associated with increased vigilance for threat-related information in general (for review, see [64]). However, trait anxiety appears to play a more specific role in dynamically regulating behavioral and neural responses to threat displays, depending on which visual pathway is engaged and on clarity or ambiguity of the threat. Our findings of the enhanced responses to averted-gaze fear (clear threat) in the high trait-anxiety individuals are largely consistent with previous work on the effect of anxiety on perception of fearful faces. For example, observing averted- gaze fear resulted in enhanced integrative processing and cuing effects for those with high (but not low) trait anxiety [11,29,55–57]. What was not clear from the previous work, however, was the differential effect of anxiety when fearful faces contained direct eye gaze (ambiguous threat cue), compared to averted eye gaze, and the contribution of the main visual pathways (M and P) to this processing. The achromatic M-pathway has characteristics that make it well suited to rapid processing of coarse, ‘gist’ information (see [65] for a review), and has been implicated in triggering top-down facilitation in object recognition in the orbitofrontal cortex [66,67], and recognition of clear threat in scene images [68,69]. Here we observed selective *facilitation* in recognizing M-biased averted-gaze fearful faces (e.g., rapid detection of clear threat cues), but also selective *impairment* in recognizing P-biased direct-gaze fearful faces (e.g., detailed analysis of ambiguous threat cues) in high trait-anxiety individuals. This finding suggests that the effect of higher trait anxiety may be to increase responsiveness to, and facilitate reflexive processing of, clear threat-related cues (e.g., averted-gaze fear). However it can be also disruptive when threat cues require detailed, reflective processing to resolve ambiguity (e.g., P-biased processing of threat ambiguity). While recognition accuracy of ambiguous facial cues (direct-gaze fear and averted-gaze neutral faces) was similar for M- and P-biased faces with low trait-anxiety, the accuracy decreased for the P-biased stimuli with anxiety.

To support and extend our behavioral findings, we also observed that high trait-anxiety individuals showed increased amygdala reactivity in a specific manner, such that the right amygdala activity increased along with anxiety only to the M-biased averted-gaze fear faces, and the left amygdala activity became greater only to the P-biased direct-gaze fear as trait anxiety increased. The effect of anxiety on amygdala attunement in the interaction of emotional valence, eye gaze direction, and pathway biasing was particularly pronounced for fearful faces but not for neutral faces, suggesting that this modulation by anxiety is specifically associated with threat detection from face stimuli. Consistent with this, we did not find any evidence for this modulation by anxiety in the fusiform face area (FFA; Supplementary Information SI2-SI4).

The existing literature on the laterality effect on emotional processing in high vs. low anxiety individuals is rather mixed. Many of the studies have shown increased right hemisphere (RH) dominance in affective processing of high-anxiety individuals. For example, a left visual field (LVF) bias has been reported for processing fearful faces in high-anxiety individuals [13] and masked angry faces presented only in LVF captured more attentional resources in high-anxiety individuals [53]. Nonclinical state anxiety was also found to be associated with increased right-hemisphere activity, as measured by regional blood flow [70], suggesting that sensitivity of right hemisphere to the presence of threat stimuli seems to be especially, although not exclusively, heightened in high-anxiety individuals. However, there are also some studies showing the association between trait anxiety and the left hemisphere (LH) activation [71,72] and larger-anxiety-related attentional bias for threatening faces presented in RVF, relative to the LVF [12,50]. Here, we directly tested for laterality effects in the left and right amygdala and found that trait anxiety modulates amygdala reactivity in both hemispheres, but with different processing emphases, such that high trait anxiety was associated with increased right amygdala reactivity to M-biased averted-gaze fear (clear threat), but increased left amygdala reactivity to P-biased direct-gaze fear (ambiguous threat). This result is also in line with the notion that reflective threat perception (e.g., resolving ambiguity from direct-gaze fear) is more left-lateralized, whereas reflexive processing (e.g., detecting clear threat from averted-gaze fear) is more right-lateralized [23,31,58,59]. Furthermore, such hemispheric lateralization became more pronounced in participants with higher trait anxiety. Thus, trait anxiety appears to play an important role in regulating the balance between the left and right amygdala responses depending on types of threat processing and pathway-biasing.

To conclude, the current study provides the first behavioral and neural evidence that trait anxiety differentially modulates the magnocellular processing of clear emotional cues (e.g., congruent combination of facial cues: averted-gaze fear and direct-gaze neutral) and parvocellular processing of ambiguous emotional cues (e.g., direct-gaze fear and averted gaze neutral). Observers’ trait anxiety also plays a specific role in differentially shaping the hemispheric lateralization in the amygdala reactivity, as a result of the complex, but systematic interplay of cue ambiguity and the visual pathway biases. Using a larger and more representative sample of a population (N=108), the current findings on the differential effects of trait anxiety on the information processing via M- vs. P-pathways and the hemispheric lateralization provide a more generalizable model for neurocognitive mechanisms underlying the perception of facial threat cues and its systematic modulation by anxiety.

## Method

### Participants

108 participants (65 female) from the Massachusetts General Hospital (MGH) and surrounding communities participated in this study. The age of the participants ranged from 18 to 70 (mean=37.05, SD=14.7). The breakdown of participants’ ethnic background is detailed in Supplementary Information SI1. All had normal or corrected-to-normal visual acuity and normal color vision, as verified by the Snellen chart [73], the Mars letter contrast sensitivity test [74], and the Ishihara color plates [75]. Informed consent was obtained from the participants in accordance with the Declaration of Helsinki. The experimental protocol was approved by the Institutional Review Board of MGH. The participants were compensated with $50 for their participation in this study.

### Apparatus and stimuli

The stimuli were generated using MATLAB (Mathworks Inc., Natick, MA), together with the Psychophysics Toolbox extensions [76,77]. The stimuli consisted of a face image presented in the center of a gray screen, subtending 5.79° × 6.78° of visual angle. We utilized a total of 24 face identities (12 female), 8 identities selected from the Pictures of Facial Affect [78], 8 identities from the NimStim Emotional Face Stimuli database [79], and the other 8 identities from the FACE database [80]. The face images displayed either a neutral or fearful expression with either a direct gaze or averted gaze, and were presented as M-biased, P-biased, or Unbiased stimuli, making 288 unique visual stimuli in the end. Faces with an averted gaze had the eyes pointing either leftward or rightward.

Each face image was first converted to a two-tone image (black-white; termed the *Unbiased* stimuli from here on). From the two-tone image, low-luminance contrast (< 5% Weber contrast), achromatic, grayscale stimuli (magnocellular-biased stimuli), and chromatically defined, isoluminant stimuli (red-green; parvocellular-biased stimuli) were generated. The low-luminance contrast images were designed to preferentially engage the M-pathway, while the isochromatic images were designed to engage the P-pathway, as such image manipulation has been employed successfully in previous studies [67,81–87]. The foreground-background luminance contrast for achromatic M-biased stimuli and the isoluminance values for chromatic P-biased stimuli vary somewhat across individual observers. Therefore, these values were established for each participant in separate test sessions, with the participant positioned in the scanner, before commencing functional scanning. This ensured that the exact viewing conditions were subsequently used during functional scanning in the main experiment. Following the procedure in Kveraga et al. [67], Thomas et al. [87], and Boshyan et al. [68], the overall stimulus brightness was kept lower for M stimuli (the average value of 115.88 on the scale of 0–255) than for P stimuli (146.06) to ensure that any processing advantages for M-biased stimuli were not due to greater overall brightness of the M stimuli, as described in detail below.

### Procedure

Before the fMRI session, participants completed the Spielberger State-Trait Anxiety Inventory (STAI; [88]). Participants were then positioned in the fMRI scanner and asked to complete the two pretests to specify the luminance values for M stimuli and chromatic values for P stimuli and the main experiment. The visual stimuli containing a face image were rear-projected onto a mirror attached to a 32-channel head coil in the fMRI scanner, located in a dimly lit room. The following procedures for pretests used to establish the isoluminance point and the appropriate luminance contrast are standard techniques and have been successfully used in many studies exploring the M- and P-pathway contributions to object and scene recognition, visual search, schizophrenia, dyslexia, and simultanagnosia [67–69,81–87].

#### Pretest 1: Measuring luminance threshold for M-biased stimuli

The appropriate luminance contrast was determined by finding the luminance threshold via a multiple staircase procedure. Figure 1A illustrates a sample trial of Pretest 1. Participants were presented with visual stimuli for 500 msec and instructed to make a key press to indicate the facial expression of the face that had been presented. They were required to choose one of the four options: 1) neutral, 2) angry, 3) fearful, or 4) did not recognize the image. One-fourth of the trials were catch trials in which the stimulus did not appear. To find the threshold for foreground-background luminance contrast, our algorithm computed the mean of the turnaround points above and below the gray background ([120 120 120] RGB value on the 8-bit scale of 0–255). From this threshold, the appropriate luminance (~3.5% Weber contrast) value was computed for the face images to be used in the low-luminance-contrast (M-biased) condition. As a result, the average foreground RGB values for M-biased stimuli were [116.5(±0.2) 116.5(±0.2) 116.5(±0.2)].

#### Pretest 2: Measuring red-green isoluminance value for P-biased stimuli

For the chromatically defined, isoluminant (P-biased) stimuli, each participant’s isoluminance point was determined using heterochromatic flicker photometry with two-tone face images displayed in rapidly alternating colors, between red and green. The alternation frequency was ~14Hz, because in our previous studies [67–69,87] we obtained the best estimates for the isoluminance point (e.g., narrow range within-subjects and low variability between-subjects; [67]) at this frequency. The isoluminance point was defined as the color values at which the flicker caused by luminance differences between red and green colors disappeared and the two alternating colors fused, making the image look steady. On each trial (Figure 1B), participants were required to report via a key press whether the stimulus appeared flickering or steady. Depending on the participant’s response, the value of the red gun in [r g b] was adjusted up or down in a pseudorandom manner for the next cycle. The average of the values in the narrow range when a participant reported a steady stimulus became the isoluminance value for the subject used in the experiment. Thus, isoluminant stimuli were defined only by chromatic contrast between foreground and background, which appeared equally bright to the observer. The average foreground red value was 151.7(±5.35) on the background with green value of 140.

#### Main experiment

Figure 1C illustrates a sample trial of the main experiment. After a variable pre-stimulus fixation period (200–400 msec), a face stimulus was presented for 1000 msec, followed by a blank screen (1100–1300 msec). Participants were required to indicate whether a face image looked fearful or neutral, as quickly as possible. Key-target mapping was counterbalanced across participants: One half of the participants pressed the left key for neutral and the right key for fearful and the other half pressed the left key for fearful and the right key for neutral. Feedback was provided on every trial.

#### fMRI data acquisition and analysis

fMRI images of brain activity were acquired using a 1.5 T scanner (Siemens Avanto) with a 32-channel head coil. High-resolution anatomical MRI data were acquired using T1-weighted images for the reconstruction of each subject’s cortical surface (TR=2300 ms, TE=2.28 ms, flip angle=8°, FoV=256x256 mm^2^, slice thickness=1 mm, sagittal orientation). The functional scans were acquired using simultaneous multislice, gradient-echo echoplanar imaging with a TR of 2500 ms, three echoes with TEs of 15 ms, 33.83 ms, and 52.66 ms, flip angle of 90°, and 58 interleaved slices (3x3x2 mm resolution). Scanning parameters were optimized by manual shimming of the gradients to fit the brain anatomy of each subject, and tilting the slice prescription anteriorly 20–30° up from the AC-PC line as described in the previous studies [67,89,90], to improve signal and minimize susceptibility artifacts in the subcortical brain regions. For each participant, the first 16 seconds of each run were discarded, followed by the actual acquisition of 96 functional volumes per run (lasting 4 minutes). There were four successive functional runs, providing the 384 functional volumes per subject in total, including the 96 null, fixation trials and the 288 stimulus trials. In our 2 (emotion: fear vs. neutral) × 2 (eye gaze direction: direct vs. averted) × 3 (bias: unbiased, M-biased, and P-biased) design, each condition had 24 repetitions, and the sequence of total 384 trials was optimized for hemodynamic response estimation efficiency using the *optseq2* software (https://surfer.nmr.mgh.harvard.edu/optseq/).

The acquired functional images were pre-processed using SPM8 (Wellcome Department of Cognitive Neurology). The functional images were corrected for differences in slice timing, realigned, corrected for movement-related artifacts, coregistered with each participant’s anatomical data, normalized to the Montreal Neurological Institute (MNI) template, and spatially smoothed using an isotropic 8-mm full width half-maximum (FWHM) Gaussian kernel. Outliers due to movement or signal from preprocessed files, using thresholds of 3 SD from the mean, 0.75 mm for translation and 0.02 radians rotation, were removed from the data sets, using the ArtRepair software [91].

Subject-specific contrasts were estimated using a fixed-effects model. These contrast images were used to obtain subject-specific estimates for each effect. For group analysis, these estimates were then entered into a second-level analysis treating participants as a random effect, using one-sample t-tests at each voxel. Age and anxiety of participants were used as covariates. In order to examine brain activations for averted and direct fear in M- and P-biased stimuli, we defined the following contrasts between conditions: M averted fear – M direct fear, M direct fear – M averted fear, P direct fear – P averted fear, and P averted fear – P direct fear. For illustration purposes, the group contrast images were overlaid onto a group average brain using MRIcroGL software (http://www.mccauslandcenter.sc.edu/mricrogl/home).

For ROI analyses, we selected a contrast between all the visual stimulation trials and baseline (e.g., Null trials). From this contrast, we used the rfxplot toolbox (http://rfxplot.sourceforge.net) for SPM and extracted the beta weights from the left and right amygdala for all the four conditions of our main interest: M-biased averted fear, M-biased direct fear, P-biased averted fear, P-biased direct fear. We used the same MNI coordinates for the left and right amygdala (x=±18, y=−2, y=16) as reported in the relevant previous work on the role of anxiety in perceiving fear faces with direct or averted eye gaze [29]. Around these coordinates, we defined 6mm spheres and extracted all the voxels from each individual participant’s functional data within those spheres. The extracted beta weights for each of the four conditions were subjected to a linear regression and correlation analyses along with the participants’ trait anxiety scores.

## Acknowledgments

This work was supported by the National Institutes of Health R01MH101194 to K.K. and to R.B.A., Jr. Kestas Kveraga: Reginald B. Adams, Jr.

## Author Contributions

R. B. Adams, and K. Kveraga developed the study concept and designed the study. Testing and data collection were performed by H. Y. Im, N. Ward, C. A. Cushing, and J. Boshyan. H. Y. Im analyzed the data and all the authors wrote the manuscript.

